# Three conserved hydrophobic residues in the CC domain of Pit contribute to its plasma membrane localization and immune induction

**DOI:** 10.1101/2021.07.31.454611

**Authors:** Qiong Wang, Yuying Li, Ken-ichi Kosami, Chaochao Liu, Jing Li, Dan Zhang, Daisuke Miki, Yoji Kawano

**Author notes:** Correspondence should be addressed to Yoji Kawano, Institute of Plant Science and Resources Okayama University 2-20-1, Chuo, Kurashiki, Okayama 710-0046, Japan. These authors contributed equally to this work (Qiong Wang, Yuying Li, Ken-ichi Kosami). **Author emails: Qiong Wang**, **Yuying Li**, **Ken-ichi Kosami**, **Chaochao Liu**, **Jing Li**, **Dan Zhang**, **Daisuke Miki:**, **Yoji Kawano**.

## Abstract

Nucleotide-binding leucine-rich repeat (NLR) proteins work as crucial intracellular immune receptors. N-terminal domains of NLRs fall into two groups, namely coiled-coil (CC) and Toll-interleukin 1 receptor (TIR) domains, which play critical roles in signal transduction and disease resistance. However, the activation mechanisms of NLRs, and how their N-termini are involved in immune induction, remain largely unknown. Here, we revealed that the rice NLR Pit self-associates through its CC domain. The CC domain of Pit possesses three conserved hydrophobic residues that are known to be involved in homodimer formation in two NLRs, barley MLA10 and Arabidopsis RPM1. Interestingly, the function of these residues in Pit is different from that in MLA10 and RPM1. Although the three hydrophobic residues are important for Pit-induced disease resistance against rice blast fungus, they do not participate in self-association or in binding to downstream signaling molecules. Based on homology modeling of Pit using the structure of the Arabidopsis NLR ZAR1, we tried to clarify the role of the three conserved hydrophobic residues and found that they are involved in the plasma membrane localization. Our findings provide novel insights for understanding the mechanisms of NLR activation as well as the relationship between subcellular localization and immune induction.

## INTRODUCTION

Plants have developed two tiers in their immune system, called pattern-triggered immunity (PTI) and effector-triggered immunity (ETI), to detect invasion by various pathogens (Jones & Dangl, 2006; Dodds & Rathjen, 2010). The initiation of PTI depends on the successful perception of conserved pathogen-associated molecular patterns (PAMPs) by surface-localized pattern recognition receptors (PRRs) (Macho & Zipfel, 2014). Once PTI signaling is activated, it is usually accompanied by a series of immune responses, such as the production of reactive oxygen species (ROS), the expression of pathogenesis-related genes, and the synthesis of antimicrobial phytoalexins and the cell wall component lignin (Bigeard *et al*., 2015). In general, PTI is sufficient to resist the attack of pathogens. Nevertheless, during evolution, pathogens have acquired the ability to secrete effectors into the apoplast or the plant cytoplasm to counteract the defense of PTI (Cui *et al*., 2015; Jones *et al*., 2016). To overcome this invasion, plants have co-evolved ETI as the second tier of the immune system. The majority of genetically characterized disease resistance traits in plants map to genes encoding nucleotide-binding domain and leucine-rich repeat proteins (NLRs). These act as receptors to surveil effectors derived from pathogens and to activate ETI, which includes the hypersensitive response (HR) and ROS production. NLRs share two core domains: a central nucleotide-binding (NB-ARC) domain and a C-terminal leucine-rich repeat (LRR) domain (Cui *et al*., 2015). The NB-ARC domain is thought to serve as a switch domain in NLRs by controlling nucleotide exchange and hydrolysis, and this nucleotide exchange leads to conformational change and oligomerization of NLRs, resulting in the triggering of ETI (Takken *et al*., 2006; Maekawa *et al*., 2011a). Highly variable LRR domains define at least part of the recognition specificity of NLRs to pathogen effector proteins. The N-terminus of NLRs is categorized into two domains, namely a Toll/interleukin-1 receptor (TIR) domain and a coiled-coil (CC) domain, and therefore NLRs are subclassified into TIR-NLRs (TNLs) and CC-NLRs (CNLs). Previous studies have demonstrated that in several NLRs, overexpression of the CC or TIR domain alone has autoactivity to induce cell death, implying that N-terminal CC and TIR domains are important platforms to trigger immune responses (Swiderski *et al*., 2009; Bernoux *et al*., 2011; Collier *et al*., 2011; Maekawa *et al*., 2011a; Wang *et al*., 2015). N-terminal CC and TIR domains are now known to play key roles in several functions, including indirect surveillance of pathogen effectors and binding of downstream signaling molecules (Cui *et al*., 2015; Jones *et al*., 2016; Kourelis & van der Hoorn, 2018).

Moreover, self (homomers) and non-self (heteromers) oligomerization of N-terminal NLRs are indispensable to trigger ETI (Maekawa *et al*., 2011a; Williams *et al*., 2014; Wang *et al*., 2019a). Structural studies have revealed that TIR domains exhibit a flavodoxin-like fold consisting of five *α*-helices surrounding a five-strand *β*-sheet, and at least two different oligomerization interfaces exist among the TIR domains (Chakraborty & Ghosh, 2020). The flax TNL L6 possesses the *α*D and *α*E helices (the DE interface) (Bernoux *et al*., 2011), and a pair of Arabidopsis TNLs, RPS4 and RRS1, have *α*A and *α*E helices (the AE interface) (Williams *et al*., 2014). Interestingly, the *Arabidopsis* TNLs SNC1 and RPP1 have the DE/AE or DE/AE-like interaction interfaces (Zhang *et al*., 2017; Chakraborty & Ghosh, 2020). Recently, two cryo-electron microscopy (EM) structures of TNLs, *Nicotiana benthamiana* ROQ1 and *Arabidopsis* RPP1, were soloved and they revealed that direct bindings of both TNLs to the corresponding effectors induce TNLs to form tetrameric resistosomes for immune signaling (Ma *et al*., 2020; Martin *et al*., 2020). The TIR domains of them bind to each other via two distinct interfaces, AE and BE (equivalent to the BB-loop of other TIR domains). While the first structure of the CC domain of CNLwas revealed as an antiparallel homodimer of barley MLA10 in crystals (Maekawa *et al*., 2011b). Subsequent studies have proved the CC domains of all the other NLRs including wheat Sr33 and potato Rx behave monomeric proteins with a four-helix bundle conformation (Hao *et al*., 2013; Casey *et al*., 2016). Recently, using a cryo-EM, Wang et al. revealed the full length structures of the Arabidopsis CNL ZAR1 in monomeric inactive and transition states as well as the active pentameric ZAR1 resistosome (Wang *et al*., 2019a). The CC domain of ZAR1 also displays a four-helix bundle conformation. A large portion of the helix *α*1 (residues 12–44) of the MLA10 CC domain appears to be an important interface for homodimerization. Single mutations in three hydrophobic residues (I33, L36, and M43) of the helix *α*1 in MLA10 dramatically decreased self-association as well as binding activity to a downstream signaling molecule, HvWRKY1, resulting in compromised resistance to the pathogenic powdery mildew fungus. The hydrophobicity of these residues is conserved among various CNLs including Arabidopsis RPM1. Triple mutation of the corresponding three hydrophobic residues in RPM1 also leads to a loss of self-association and immune induction activity (El Kasmi *et al*., 2017). All of the single mutants in three hydrophobic residues show reduced interaction with the small host protein RIN4 (El Kasmi *et al*., 2017). It is unclear whether these residues are universally involved in self-association or whether their functions differ in each NLR.

Evidence has been accumulating that the subcellular distribution of NLRs is important for their functions. Perception of the fungal effector A10 by MLA10 triggers the nuclear translocation from the cytosol, resulting in interaction between MLA10 and HvWRKY1 in the nucleus to induce defense responses (Shen *et al*., 2007). The intranuclear activities of the Arabidopsis TNL RPS4 restrict bacterial growth and programmed cell death, and nucleocytoplasmic coordination of RPS4 needs transcriptional resistance reinforcement (Heidrich *et al*., 2011). The potato CNL Rx1 is located in both the cytoplasm and the nucleus, and the appropriate nucleocytoplasmic distribution of Rx1 is required for full functionality (Slootweg *et al*., 2010; Tameling *et al*., 2010). Rx1 activation triggered by effector recognition occurs only in the cytoplasm. The CC domain and the cochaperone SGT contribute to nuclear localization of Rx1, but the LRR domain is associated with cytoplasmic localization. The Arabidopsis CNL RPM1 requires plasma membrane distribution while the potato CNL R3a needs endomembrane localization, and disrupting the proper localization of both NLRs impairs their functions (Gao *et al*., 2011; Engelhardt *et al*., 2012). The oligomerization-induced active Arabidopsis ZAR1 complex associates with the plasma membrane (Wang *et al*., 2019a). Although our knowledge of NLR protein localization has increased in recent years, it is not yet sufficient to understand the mechanisms and significance of the dynamic nature of NLR protein localization or the relationship between subcellular localization and activation states.

We have previously revealed that the small GTPase OsRac1 functions as a molecular switch in rice and plays key roles in both PTI and ETI (Kawano *et al*., 2010; Akamatsu *et al*., 2013; Kawano & Shimamoto, 2013; Kawano *et al*., 2014b). OsRac1 forms immune protein complex(es) directly or indirectly with 16 binding partners such as NADPH oxidase and OsMPK6, thereby leading to the induction of immune responses (Lieberherr *et al*., 2005; Chen *et al*., 2010; Akamatsu *et al*., 2013; Kawano & Shimamoto, 2013; Kawano *et al*., 2014b; Kosami *et al*., 2014). OsRac1 acts as a downstream switch molecule for three CNLs, Pit, Pia, and PID3, which all confer resistance to *Magnaporthe oryzae*, implying that OsRac1 is a key signaling switch for rice CNLs (Ono *et al*., 2001; Kawano *et al*., 2010; Wang *et al*., 2018; Zhou *et al*., 2019). Recently, we clarified how the CNL Pit activates OsRac1. Pit interacts directly with the GDP/GTP exchanger (GEF) protein OsSPK1, which is an activator for OsRac1, through its CC domain (Wang *et al*., 2018) and also associates with OsRac1 through its NB-ARC domain (Kawano *et al*., 2010). Both Pit and OsRac1 seem to be posttranslationally modified by a lipid modification, palmitoylation, and these three proteins may form a ternary complex at the plasma membrane to trigger ETI (Ono *et al*., 2001; Kawano *et al*., 2014a; Yalovsky, 2015; Wang *et al*., 2018).

In this study, we clarified the role of the above-mentioned three conserved hydrophobic residues in the CC domain of Pit. Interestingly, the three residues are involved in the plasma membrane localization of Pit, and are indispensable for Pit-mediated disease resistance to rice blast fungus, but do not participate in self-association and binding to its direct signaling molecules OsSPK1 and OsRac1. Collectively, our results shed light on how NLRs trigger immune induction.

## RESULTS

### Pit self-associates through its CC domain

Since several CNL and TNL proteins have been reported to self-associate (Mestre & Baulcombe, 2006; Ade *et al*., 2007), we tested whether the rice NLR Pit forms homo-oligomers *in planta*. We transiently co-expressed full-length Pit WT-HA and Pit WT-Myc in *Nicotiana benthamiana* and performed a co-immunoprecipitation (co-IP) assay. When Pit WT-HA was precipitated with anti-HA antibody, Pit WT-Myc co-precipitated but a control GUS-HA did not, indicating that Pit self-associates *in planta* (Figure 1A). Previous studies have demonstrated that the N-terminal CC and TIR domains are important interfaces for homo-oligomerization in NLRs, and that these interactions are indispensable for NLR functions (Maekawa *et al*., 2011a; Williams *et al*., 2014). Next, we examined whether Pit self-associates through its CC domain (amino acids 1-140). By using a yeast two-hybrid assay, we found that the CC domain of Pit formed homomers (Figure 1B). Consistent with this observation, self-association between the CC domains of Pit was observed in a co-IP assay in *N. benthamiana* (Figure 1C) and an *in vitro* binding assay (Figure 1D). Taken together, these results indicate that Pit forms homomers through its CC domain.

**Figure 1.**
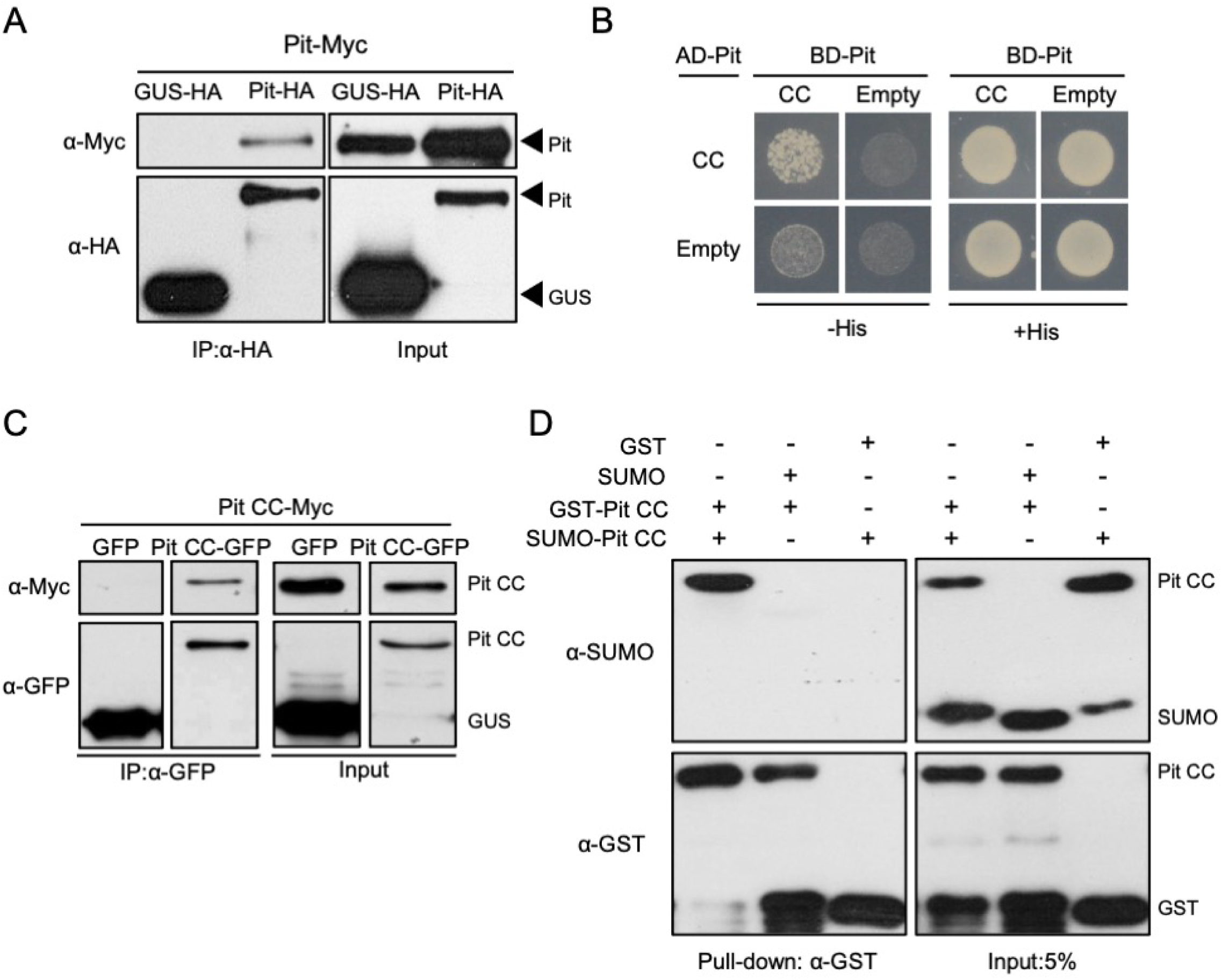
The CC domain and full-length Pit self-associate. **A**, Co-IP assay to assess self-association of full-length Pit in *N. benthamiana*. Total protein extract was immunoprecipitated with anti-HA antibody, and western blotting was then carried out with anti-HA and anti-Myc antibodies. **B**, Yeast two-hybrid assay to test self-association of the Pit CC domain. Growth of yeast cells coexpressing GAL4-AD or GAL4-BD fused with the CC domain of Pit on selective medium without histidine (-His) represents a positive interaction. AD: GAL4 activation domain, BD: GAL4 DNA-binding domain. **C**, Co-IP assay to assess self-association of the Pit CC domain in *N. benthamiana*. Total protein extract was immunoprecipitated with anti-GFP antibody, and western blotting was then carried out with anti-GFP and anti-Myc antibodies. **D**, GST pull-down assay to verify the self-association of the Pit CC domain. Purified GST or GST-tagged Pit CC immobilized on Sepharose was incubated with His- SUMO-tagged Pit CC. After washing, the bound proteins were eluted by addition of SDS loading buffer for immunoblotting with anti-GST and anti-SUMO.

### Three conserved residues do not contribute to self-association of Pit

Although the primary sequences of the N-terminal CC domain of NLRs are dissimilar, three hydrophobic residues (I33, L36, and M43) of the helix *α*1 in MLA10 are highly conserved among the known CNL proteins including Pit (Maekawa *et al*., 2011a) (Figure 2A). In MLA10 and RPM1, these three residues are involved in self-association and are indispensable for immune induction (Maekawa *et al*., 2011b; El Kasmi *et al*., 2017). I33, L36, and M43 in MLA10 correspond to I34, L37, and L41 in Pit. Our finding that Pit forms homomers through its CC domain raised the possibility that the three conserved hydrophobic residues of Pit also participate in oligomerization and are essential for its function. To test this hypothesis, we built a homology model of the CC domain of Pit using the crystal structure of the CC domain of MLA10 (Maekawa *et al*., 2011a). Similar to MLA10, the model structure of the CC domain of Pit dimerized through the helix *α*1 using three hydrophobic residues of I34, L37, and L41 (Figure 2B). We generated CC domain Pit mutants in which these three conserved residues were converted to negatively charged glutamic acid, and tested whether they are involved in self-association. Interestingly, both the single mutations (Pit I34, L37, or L41) and the triple mutation of Pit (Pit 3E: Pit I34E L37E L41E) retained self-association ability in an *in vitro* binding assay (Figure 2C), and a consistent result was obtained in a yeast two-hybrid assay (Figures 2D and S1A). We also conducted a co-IP assay in *N. benthamiana* using full-length Pit but could not observe a visible effect on self-association (Figure 2E). Overall, these results indicate that the three hydrophobic residues do not contribute to self-association of Pit.

**Figure 2.**
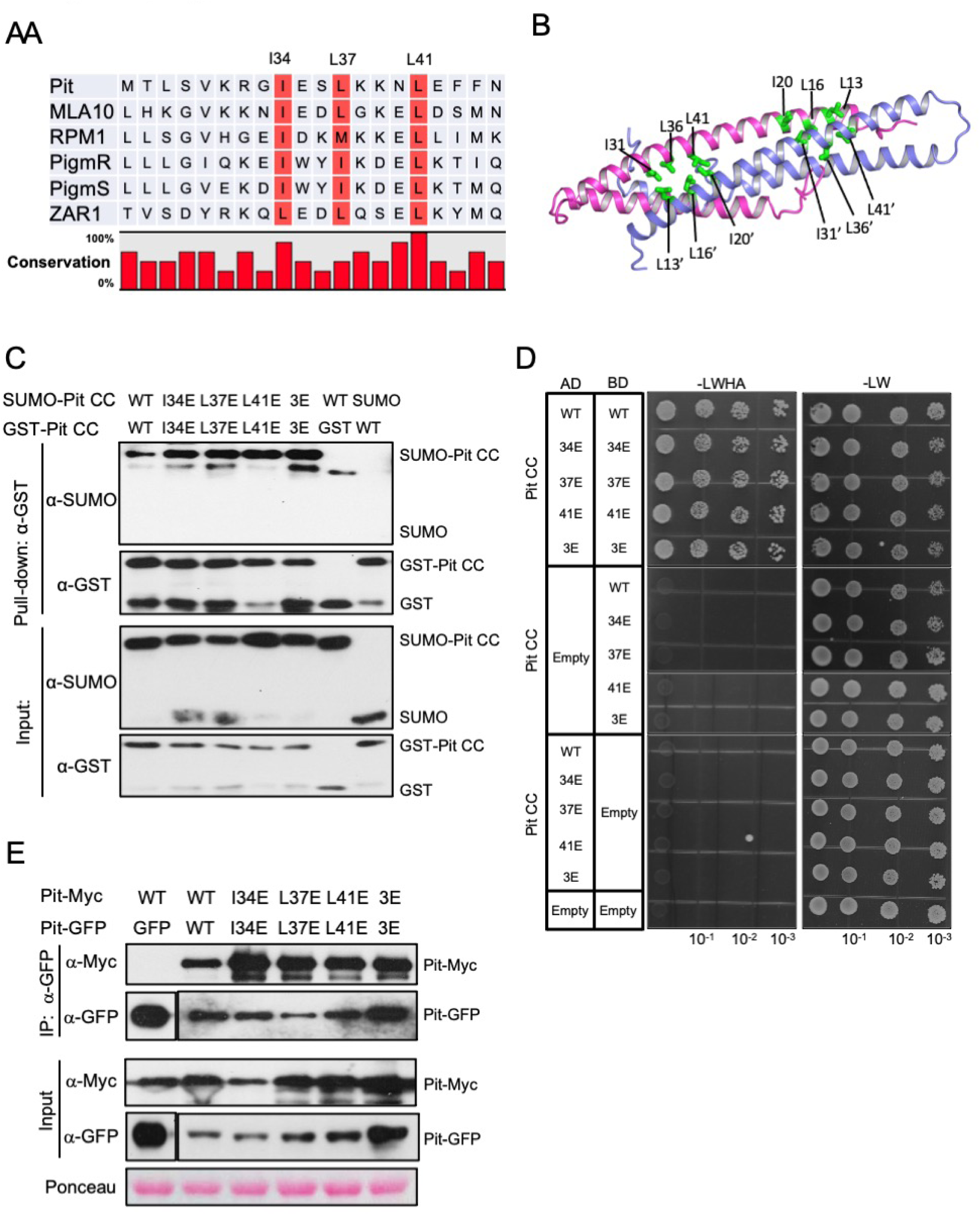
Conserved hydrophobic residues in the Pit CC domain are not involved in Pit self-association. **A**, Multiple alignment of Pit with various CNLs. **B**, Model structure of the Pit CC domain, based on the MLA10 CC domain (Protein Data Bank ID code 3QFL), shows the elongated dimer (blue and pink), stabilized by hydrophobic residues (I34, L37, and L41). The figure was drawn using PyMOL. **C**, *In vitro* pull-down assay to test the self-association of Pit CC mutants. Purified GST or GST-tagged Pit CC mutants immobilized on Sepharose was incubated with His-SUMO-tagged Pit CC mutants. After washing, the bound proteins were eluted by addition of SDS loading buffer for immunoblotting. Anti-GST and anti-SUMO antibodies were used for western blotting. **D**, Yeast two-hybrid assay to test self-association of Pit CC mutants. Growth of yeast cells coexpressing GAL4-AD or GAL4-BD fused with the CC domain of Pit on selective medium (-LWHA) represents a positive interaction. 10^-1^, 10^-2^, and 10^-3^ indicate dilution ratio. **E**, Co-IP assay to examine self-association of full-length Pit mutants in *N. benthamiana*. Total protein extract was immunoprecipitated with anti-GFP antibody, and western blottingwas then carried out with anti-GFP and anti-Myc antibodies. The post-transfer membrane was stained with Ponceau S.

### Mutations in the three conserved residues of Pit compromise Pit-mediated immune responses

Next, we examined whether the effects of the hydrophobic residue mutants of Pit influenced immune responses. We have previously generated a constitutively active form of Pit, named Pit D485V. Pit D485V is a MHD motif mutant that is able to induce cell death and ROS production in *N. Benthamiana*, probably through the employment of tobacco orthologs of OsRac1 and OsSPK1 as downstream signal transducers because the overexpression of the dominat negative form of OsRac1 suppresses Pit D485V- induced cell death in *N. Benthamiana* (Kawano *et al*., 2010; Kawano *et al*., 2014b). The single mutations and the triple mutation in the three hydrophobic residues clearly attenuated Pit D485V-induced cell death (Figures 3A and S1C) and ROS production (Figures 3B, S1B and S1C). We also employed two rice systems to evaluate the Pit mutants. We used a luciferase reporter system to monitor the effect of the Pit mutants on cell death in rice protoplasts. In this system, we transfected the *Pit* mutants with a *luciferase* vector into rice protoplasts and measured the viability of protoplasts based on luminescence. We found that the luciferase activity in cells expressing Pit WT was significantly lower than that in cells expressing control GUS, indicating that Pit WT is autoactive and induces cell death in rice protoplasts (Figure 3C). This Pit WT-induced cell death was abolished by the introduction of the mutations in three conserved hydrophobic residues (Figures 3C and S1D). Next, we tested the effect of the hydrophobic residue mutants of Pit on disease resistance to rice blast fungus. In this experiment, we generated transgenic plants of the susceptible rice cultivar Nipponbare carrying the exogenous *Pit* resistance genes (Figure S2A), and chose the avirulent rice blast fungus *M. oryzae* race 007.0, because Pit-dependent disease resistance has been established between *Pit* and *M. oryzae* race 007.0 (Hayashi *et al*., 2010). Therefore, Nipponbare is a suitable cultivar to assess transgenes encoding the *Pit* mutants. Nipponbare expressing *Pit* WT displayed shorter lesions induced by *M. oryzae* than did Nipponbare, but this effect was compromised in both single and triple mutants for the three hydrophobic residues (Figures 3D and 3E). We also precisely quantified fungal invasion by measuring the amount of fungal DNA using real-time PCR (Figure 3F). The result of this qPCR was consistent with that of the lesion length comparison, showing that mutation of the three hydrophobic residues perturbed Pit-triggered resistance to avirulent rice blast fungus. Take together, these data indicate that the three hydrophobic residues in the CC domain of Pit are indispensable for Pit-mediated immune responses.

**Figure 3.**
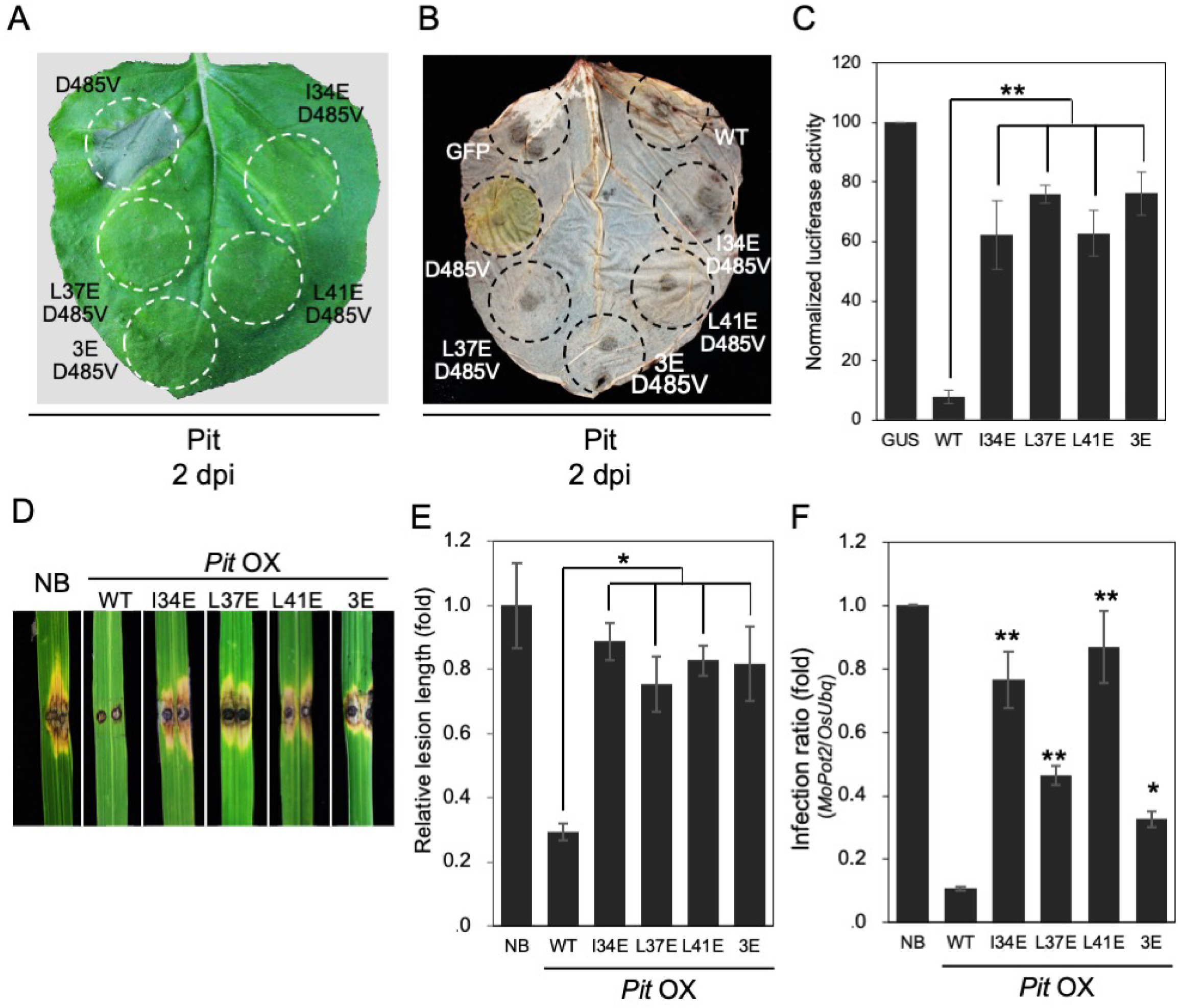
Conserved hydrophobic residues in the Pit CC domain contribute to Pit-mediated immune signaling. **A**, Cell death phenotypes induced by transient expression of Pit mutants in *N. benthamiana*. Photos were taken at 2 dpi. The circles indicate the infiltrated regions. **B**, Effect of three hydrophobic residues on Pit D485V-induced ROS production in *N. benthamiana*. ROS production was examined by DAB staining at 2 dpi. **C**, Cell death activity of Pit mutants in rice protoplasts. Relative luciferase activity (GUS=100) is shown. Data are expressed as mean ± standard error (SE) (\*\**P* < 0.01, *n* = 3). **D** and **E**, Responses, in plants overexpressing Pit WT or mutants, to infection with the incompatible *M. oryzae* race 007.0. **D**, Photograph shows typical phenotypes of transgenic and WT plants at 7 dpi. **E**, Statistical analysis of lesion length was performed at 6 dpi. Relative lesion length [Nippobnare (NB) = 1] is shown. Data are expressed as mean ± standard error (SE) (\**P* < 0.05; *n* ≥ 30). **F**, Growth of the incompatible *M. oryzae* race in Nipponbare wild-type plants and transgenic plants overexpressing Pit WT or mutants. Relative infection ratio (NB = 1) is shown. Data are expressed as mean ± standard error (SE) (\**P* < 0.05; \*\**P* < 0.01; *n* = 10).

### Mutations in the three conserved hydrophobic residues abolish Pit-induced OsRac1 activation but do not affect the interaction with OsRac1 and OsSPK1

Since the three hydrophobic residue mutations of Pit significantly perturbed Pit-mediated immune responses (Figure 3), we checked the interactions between Pit and its two downstream signaling molecules: the molecular switch of rice immunity OsRac1 and its activator OsSPK1. Pit may form a ternary complex with OsSPK1 and OsRac1 at the plasma membrane and activates OsRac1 through OsSPK1 to induce Pit-mediated immunity (Kawano *et al*., 2010; Wang *et al*., 2018). We previously mapped the binding region of OsSKP1 in Pit and revealed that a proline-rich motif of the CC domain in Pit (residues 91–95) is required for its binding to OsSPK1 (Wang *et al*., 2018). Consistent with that finding, there is no visible effect in any of the three hydrophobic residue mutants of Pit on binding to OsSPK1 in a co-IP assay in *N. benthamiana* using the CC domain (Figure 4A) and an *in vitro* binding assay (Figure S2B) and the full-length polypeptide (Figure S2C) of Pit. Moreover, mutating the three hydrophobic residues of Pit did not change its binding activity to OsRac1, probably because OsRac1 binds to the NB-ARC domain of Pit (Figure 4B) (Kawano *et al*., 2010). Next, we checked OsRac1 activation by the Pit mutants using a Förster resonance energy transfer (FRET) sensor called Ras and interacting protein chimeric unit (Raichu)-OsRac1 (Wong *et al*., 2018). In this sensor, intramolecular binding of the active GTP-OsRac1 to CRIB brings CFP closer to Venus, enabling FRET from CFP to Venus when OsRac1 is activated (Wong *et al*., 2018). The resulting Venus fluorescence represents the activation state of OsRac1 *in vivo*: low and high ratios of Venus/CFP fluorescence correspond to low and high levels of OsRac1 activation, respectively. The ratio of Venus/CFP fluorescence of Raichu-OsRac1 in rice protoplasts expressing Pit D485V was much higher than that in protoplasts expressing a control GUS, indicating that Pit D485V activates OsRac1 in rice protoplasts, but the triple mutant Pit 3E with the D485V mutation failed to trigger this activity (Figures 4C and 4D). Thus, we conclude that Pit 3E retains binding activity to OsSPK1 and OsRac1 but loses the ability of wild-type Pit to trigger OsRac1 activation.

**Figure 4.**
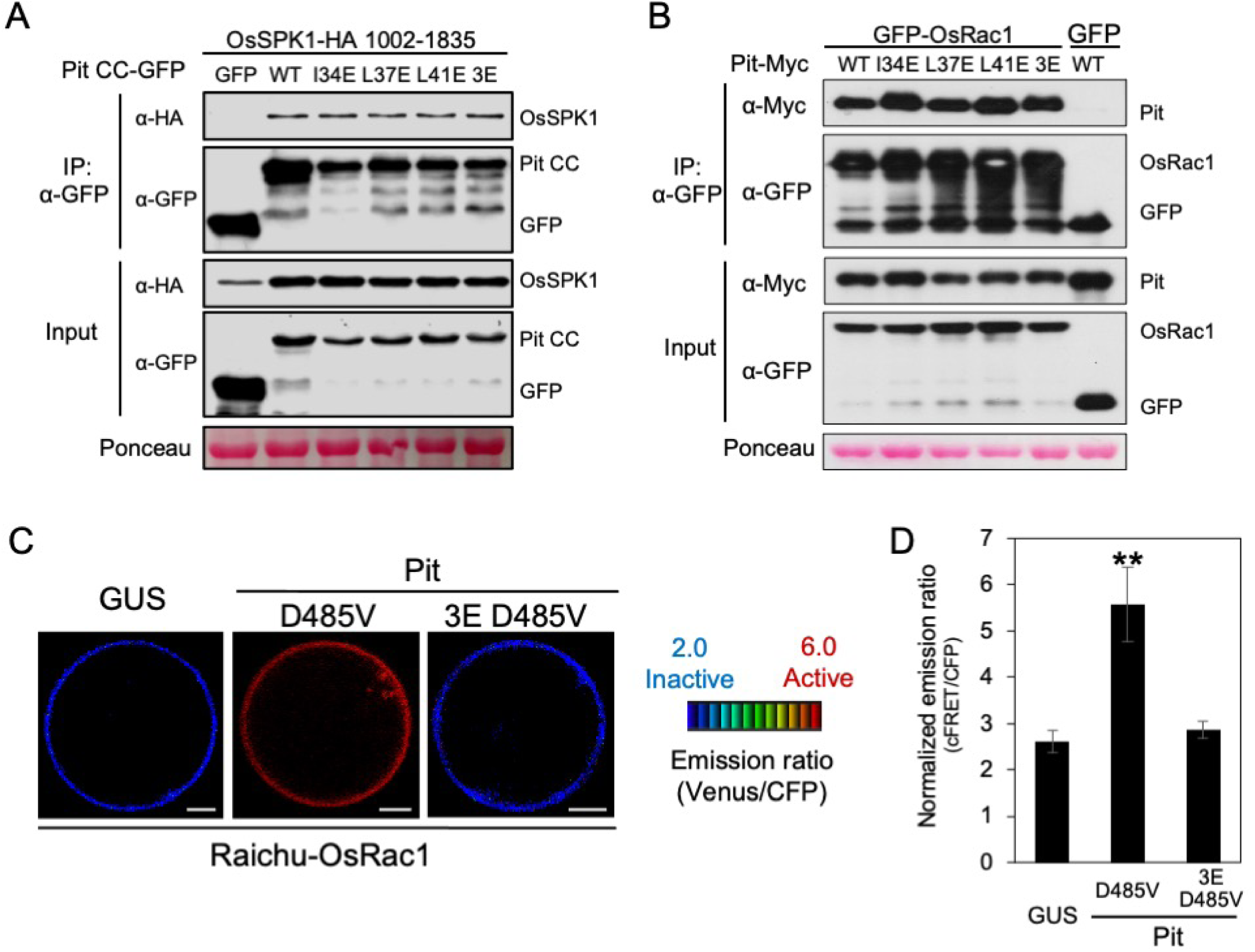
Mutations in three hydrophobic residues do not affect binding to OsSPK1 or OsRac1 but perturb Pit-mediated OsRac1 activation. **A**, Co-IP to test the interaction between OsSPK1 and Pit hydrophobic residue mutants in *N. benthamiana*. Total protein extract was immunoprecipitated with anti-GFP antibody, and western blotting was then carried out with anti-GFP and anti-HA antibodies. The post-transfer membrane was stained with Ponceau S. **B**, Co-IP to test the interaction between OsRac1 and Pit hydrophobic residue mutants in *N. benthamiana*. Total protein extract was immunoprecipitated with anti-GFP antibody, and western blotting was then carried out with anti-GFP and anti-Myc antibodies. The post-transfer membrane was stained with Ponceau S. **C** and **D**, *In vivo* OsRac1 activation by Pit hydrophobic residues mutants. **C**, Emission ratio images of confocal laser-scanning micrographs of rice protoplasts coexpressing Raichu-OsRac1 and the indicated Pit mutants, or negative control GUS. Scale bars, 5 μm. **D**, Quantification of normalized emission ratios of Venus to CFP. Data are expressed as mean ± standard error (SE) (\*\**P* < 0.01, *n* = 60).

### Homology modeling of Pit

From the results of our interaction studies in Figure 2, it appears that the MLA10 structure is not applicable to Pit. The structures of the CC domain of NLRs resolve into two types: MLA10 forms dimers and shows a helix–loop–helix structure (Maekawa *et al*., 2011a), while Sr33 and Rx display a distinct structure that exhibits a four-helix bundle (Hao *et al*., 2013; Casey *et al*., 2016). Recently, Wang et al. solved the structure of the inactive and active states of the full-length CNL ZAR1, which revealed that the N-terminal CC domain of inactivated ZAR1 (PDB code 6J5W, Chain A, 1–113) possesses a four-helix bundle, like Sr33 (PDB code 2NCG) and Rx (PDB code 4M70, Chain A) (Figure 5A), implying that Pit also displays the four-helix bundle (Hao *et al*., 2013; Casey *et al*., 2016; Wang *et al*., 2019b). To test this hypothesis and understand the function of I34, L37, and L41 in Pit, we undertook detailed homology structure modeling of Pit based on the inactive (ADP-bound) and active (dATP-bound) structures of the NLR ZAR1 (Wang *et al*., 2019a; Wang *et al*., 2019b). The model structure of the CC domain of Pit displays a four-helix bundle, and the three hydrophobic residues are buried inside the CC domain (Figure 5B). These residues locate on α-helix 2 (α2) and make hydrophobic contact with α-helix1 (α1) and α-helix 3 (α3), which may enhance the stability of the four-helix bundle. The three hydrophobic residues are conserved in Sr33 (I33, L36, and L40), Rx (L24, F27, and L31), and ZAR1 (L31, L34, and L38), and they also form similar hydrophobic contacts (Casey *et al*., 2016; El Kasmi *et al*., 2017) (Figure 5B). In the model structure of the inactive form of Pit, the LRR domain sequesters Pit in a monomeric state (Figure 5C). The CC domain of Pit contacts the helical domain (HD1) and a winged-helix domain (WHD) in the NB-ARC domain, and these interactions may keep the CC domain inactive (Burdett *et al*., 2019; Wang *et al*., 2019b).

**Figure 5.**
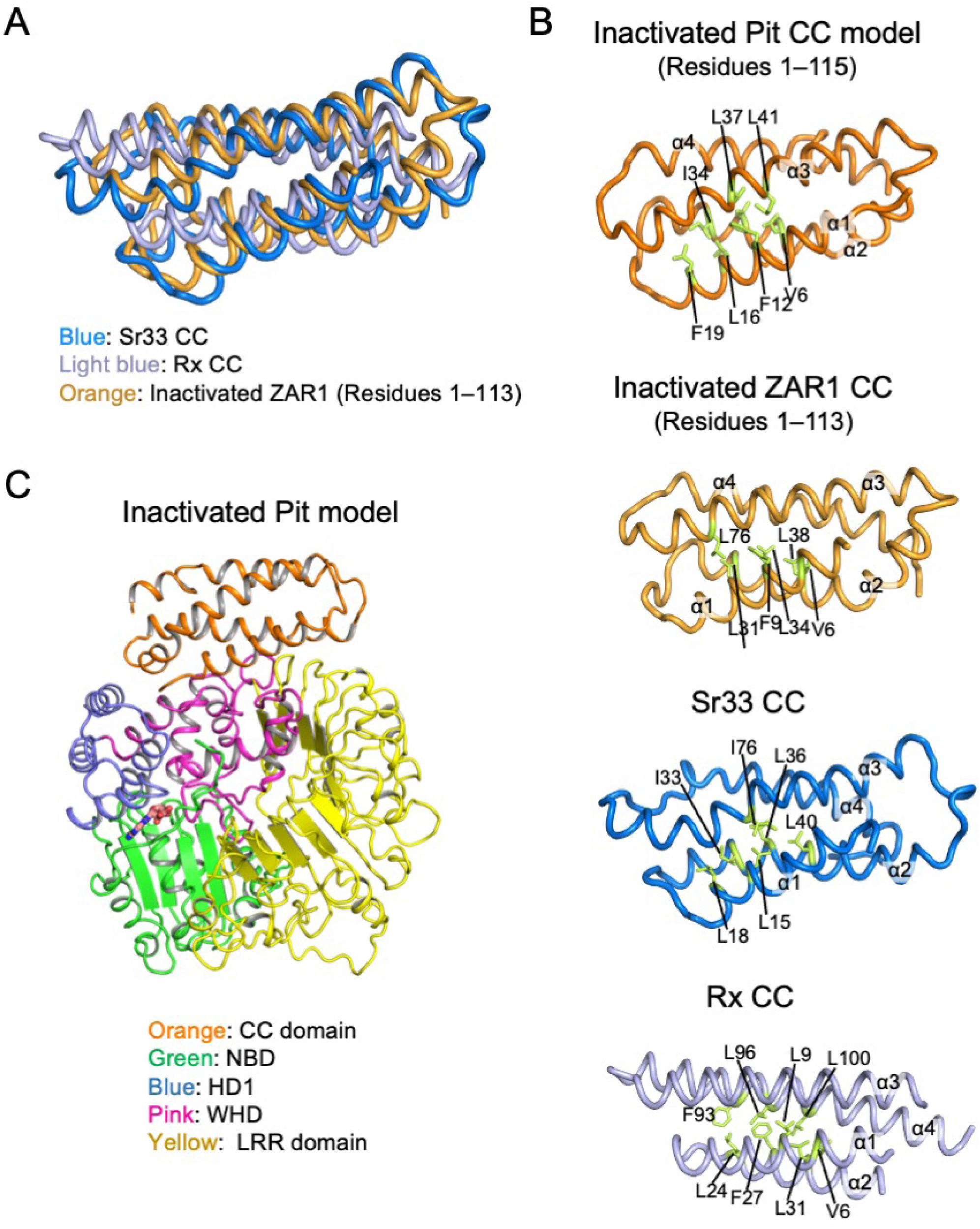
The CC domain of various NLRs. **A,** The main chains of the CC domain structure of Sr33 (blue, solution NMR condition, Protein Data Bank ID code 2NCG), Rx (light blue, crystal condition, Protein Data Bank ID code 4M70), and inactivated ZAR1 (orange, electron microscopy condition, 6J5W, residuces 1–113) were superimposed using PyMOL. **B,** Comparison of conserved hydrophobic residues of Pit (residuces 1–115), inactivated ZAR1 (residuces 1–113), Sr33, and Rx. The side chains of three key hydrophobic residues in Pit (I34, L37, and L41) and equivalent residues in inactivated ZAR1 1–113 (L31, L34, and L38), Rx (I33, L36, and L40), Sr33 (L24, F27, and L31), as well as the side chains of amino acids thought to be involved in hydrophobic interactions with these three residues, are shown in stick representation. **C,** Model structure and domain composition of full-length Pit, based on inactivated ZAR1 (Protein Data Bank ID code 6J5W), shows the monomeric state. The figure was drawn using PyMOL.

ZAR1 transitions from a monomeric inactive form to the active form, a wheel-like pentameric resistosome, during immune activation (Wang *et al*., 2019a; Wang *et al*., 2019b). Since Pit forms homomers (Figure 1), we also generated a model structure of Pit with reference to the structure of the active form of ZAR1 (Wang *et al*., 2019a). Superposition of the inactive Pit model structure with one protomer of the active Pit model structure revealed that the conformational change between the active and inactive forms of Pit probably occurs at two regions: around the hinge linking the HD and WHD domains, and in the α1 helix of the CC domain (Figure 6A). In the inactive Pit model structure, the amphipathic α1 helix is buried in the four-helix bundle and interacts with the WHD and LRR domains. In the active Pit model structure, the α1 helix rotates and separates from the four-helix bundle, becoming a fully solvent-exposed α1 helix (Burdett *et al*., 2019). In the active-form Pit model structure, Pit forms a wheel-like pentamer and all the subdomains of Pit are involved in this oligomerization (Figure 6B). The formation of an α-helical funnel-shaped structure in the CC domain contributes to the oligomerization of Pit and is consistent with the self-association of Pit through its CC domain (Figures 1 and 6B) (Wang *et al*., 2019a). In the active Pit model structure, I34, L37, and L41 locate on the α2 helix and are buried inside the CC domain, implying that they do not contribute to self-association of the CC domain. This model structure fits well with the result of our binding assays (Figure 2). Interestingly, the three conserved hydrophobic residues make hydrophobic contacts with V75, I78, and V79 of the α3 helix, which itself forms hydrophobic interactions with isoleucines I500 and L510 in the WHD domain (Figure 6C). The introduction of negatively charged glutamic acid into I34, L37, and L41 of Pit may decrease the molecular packing density between α2 and α3 helices in the CC domain, leading to a weakening of the hydrophobic interactions among the α2 and α3 helices of the CC domain, and the WHD domain. Besides, D77 in the α3 helix also forms a hydrogen bond with K532 in the LRR domain. These interactions appear to provide a foundation when the activated protein oligomerizes via its CC domain to form a functional funnel-shaped structure. We checked whether ZAR1 has these interactions between the CC, WHD, and LRR domains in the active ZAR1 structure, and found that L31 and L34 of the α2 helix make hydrophobic contacts with I75 and L76 of the α3 helix, and L115 and I118 of the α4 helix (Figure 6D). Moreover, hydrogen bonds were predicted to between the α2 helix and WHD (K46 and S413) and between the α3 helix and the LRR domain (E67 and R513, E73 and R533). These structural features of ZAR1 are similar to those of Pit modeling.

**Figure 6.**
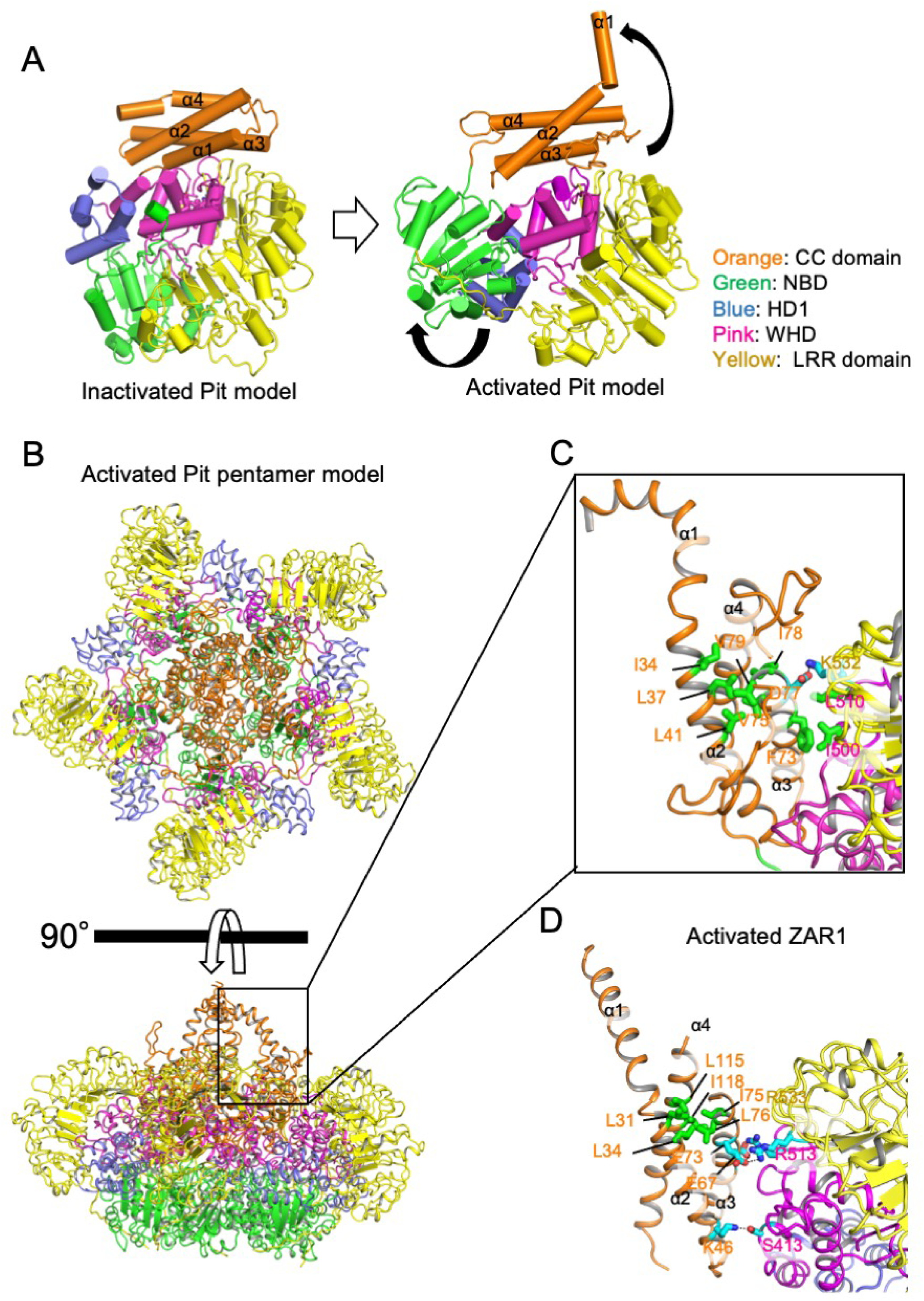
Homology modeling of Pit using activated ZAR1 as a template. **A,** Comparison of the inactivated and activated Pit model structures. The model structures of inactivated and activated Pit are based on the structures of inactivated ZAR1 (Protein Data Bank ID code 6J5W) and activated ZAR1 (Protein Data Bank ID code 6J5T). Conformational changes (black arrows) between the activated and inactivated forms of Pit occur around the hinge linking the HD domain (blue) and WHD domain (pink) of Pit and also at the α1 helix of the CC domain (orange). **B,** Model structure of the activated Pit pentamer. The extreme N-terminal α1 helix of the Pit pentamer may be required for the plasma membrane association of Pit. The CC, NBD, HD1, WHD, and LRR domains are shown in orange, green, blue, pink, and yellow, respectively. **C,** Hydrophobic interactions among α2 (I34, L37, and L41), α3 (F73, V75, I78, and V79), and WHD domain (I500 and L510) in the activated Pit pentamer model structure are shown. Residues that may be important for hydrophobic interactions in Pit function are shown in green, and hydrogen-bonded side chains are shown in light blue. **D,** Comparison of the interaction around α1 of the activated ZAR1 structure (Protein Data Bank ID code 6J5T) and the active Pit oligomer model structure. Residues involved in hydrophobic interaction around α1 are shown in green, and hydrogen-bonded side chains are shown in light blue. The figure was drawn using PyMOL.

### Mutations in the three hydrophobic residues of Pit perturb its plasma membrane localization

Next, we checked the localization of the hydrophobic residue mutants of Pit in rice protoplasts. We had previously demonstrated that Pit WT is localized at the plasma membrane, but we now found that introducing single mutations into the three hydrophobic residues compromised Pit’s plasma membrane localization (Figure 7A) (Kawano *et al*., 2010; Kawano *et al*., 2014a). We further investigated the localization of the hydrophobic residue mutants in *N. benthamiana* and found that Pit WT was well merged with FM4-64, a plasma membrane marker, confirming that Pit WT is localized in the plasma membrane (Figure 7B); in contrast, plasma membrane localization was disrupted in all of the hydrophobic residue mutants. Taken together, these results indicate that the three conserved hydrophobic residues of Pit are required for its proper plasma membrane localization.

**Figure 7.**
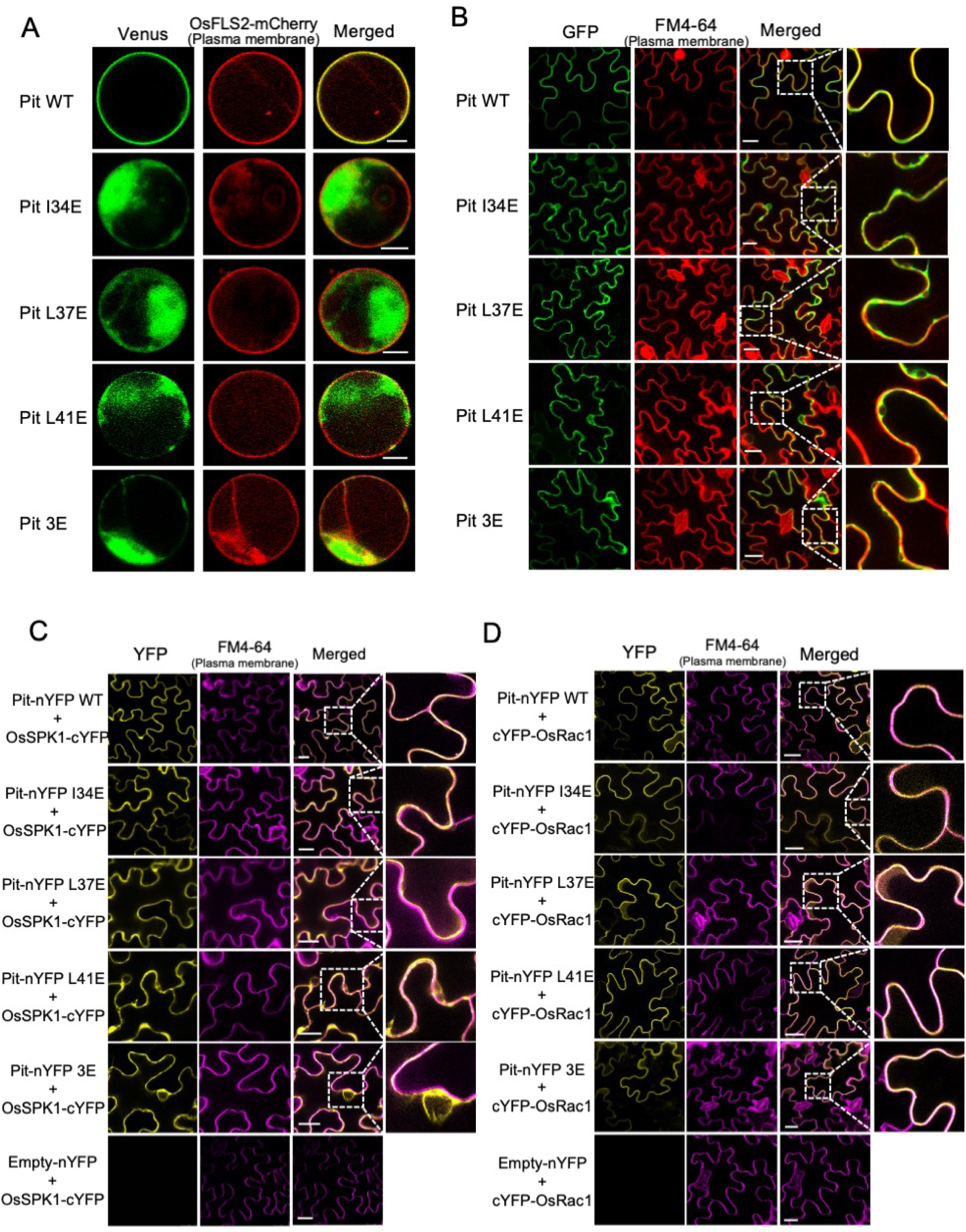
Mutations in the conserved hydrophobic residues of Pit influence its plasma membrane localization. **A** and **B**, Subcellular localization of Pit mutants in rice protoplasts and *N. benthamiana* leaves. **A**, Rice protoplasts were cotransfected with the indicated *Pit-Venus* mutants and *OsFLS2-mCherry.* Scale bars, 5 μm. **B**, Tobacco leaves were injected with *Agrobacterium* carrying *Pit-GFP* mutants (green) and stained with FM4-64 (red: plasma membrane marker). Enlarged images of the boxed areas are shown in the right panels. Scale bars, 25 μm. **C** and **D**, BiFC to detect interactions between Pit hydrophobic residue mutants and OsSPK1 (**C**) or OsRac1 (**D**). Expression constructs were transiently expressed in *N. benthamiana* after agroinfiltration. Empty vector served as a negative control. FM4-64 was used as a plasma membrane marker. Images were captured at 45 h post-infiltration. Enlarged images of the boxed areas in (C) are shown in the right panels. Scale bars, 25 μm.

We checked the OsSPK1-binding activity of these Pit mutants by bimolecular fluorescence complementation (BiFC) assay in *N. benthamiana*. Consistent with the results of the binding assays (Figure 4A and 4B), OsSPK1 binding was comparable in the mutants to that in Pit WT (Figure 7C). However, the localization of the Pit–OsSPK1 complex differed between Pit WT and the hydrophobic mutants. Pit WT interacted with OsSPK1 at the plasma membrane, as reported previously (Figure 7C) (Wang *et al*., 2018), but a large proportion of the complexes between OsSPK1 and the Pit mutants was mislocalized away from the plasma membrane. Finally, we examined complex formation between OsRac1 and the Pit mutants by a BiFC assay and found that the Pit WT-OsRac1 complex was situated at the plasma membrane. Interestingly, none of the mutations in the hydrophobic residues disrupted the Pit-OsRac1 interaction at the plasma membrane (Figure 7D), probably because OsRac1 is anchored there by its lipid modification. Taken together, these results indicate that the three conserved hydrophobic residues of Pit are required for its plasma membrane localization.

## DISCUSSION

Several TNLs and CNLs have been reported to self-associate through their N-terminal CC or TIR domains; hence, self-association via their N-termini appears to be a general feature of NLRs (Ade *et al*., 2007; Maekawa *et al*., 2011a; El Kasmi *et al*., 2017). Here, we found that the rice blast resistance protein Pit also self-associates through at least its CC domain (Figure 1). Full-length Pit forms homomers in the absence of a pathogen effector, suggesting that it may self-associate before activation and behave like other NLRs, such as RPM1, RPS5, and MLA (Ade *et al*., 2007; El Kasmi *et al*., 2017). Previous biophysical analyses have shown that MLA10 is a monomer in solutions but has a dimeric helix-loop-helix structure in crystals (Maekawa *et al*., 2011a; Casey *et al*., 2016). It is possible that the dimeric helix-loop-helix structure of MLA10 occurs under the special condition because MLA10 is predominantly monomeric in solution and its character in solution is different from that in the crystals (Casey *et al*., 2016; Bentham *et al*., 2018; Burdett *et al*., 2019). Moreover, the CC domain structures of all other NLRs, including Rx, Sr33, and ZAR1, exhibit a four-helix bundle structure (Casey *et al*., 2016; El Kasmi *et al*., 2017; Wang *et al*., 2019a). Single mutations in the hydrophobic residues (I33, L36, and M43) of α1 helix of MLA10 markedly suppressed self-association, resulting in compromised resistance to *Blumeria graminis* f. sp. *hordei*. We found that introducing the single and triple mutations into Pit did not affect homomer formation, indicating that these residues make at most a marginal contribution to self-association (Figure 2). It is possible that Pit displays a four-helix bundle structure, similar to other NLRs such as Rx, Sr33, and ZAR1. We attempted to clarify the structure of the CC domain of Pit and produced an expression system for the CC domain and full-length Pit protein using *E. coli* and insect cells, but we were unable to obtain intact Pit proteins due to difficulties in expression. The structural analysis of the Pit CC domain will be a topic for future research.

Several CNLs, including RPM1 (Gao *et al*., 2011), RPS2 (Axtell & Staskawicz, 2003), RPS5 (Qi *et al*., 2012), and Tm-2^2^ (Chen *et al*., 2017), have been reported to be localized in the plasma membrane, and this localization is indispensable for their immune induction. RPM1 appears to anchor to the plasma membrane through the plant guardee protein RIN4, which is localized to the membrane via palmitoylation (Kim *et al*., 2005). Two lipid modifications, myristoylation, and palmitoylation, in the CC domain of RPS5, participate in its plasma localization, protein stability, and function in an additive manner. We have previously revealed that a free N-terminus of Pit is required for its function because the N-terminal fusion of GFP compromises cell death activity (Kawano *et al*., 2014a). Consistent with this, Pit has two palmitoylation sites in its CC domain, which play a key role in the plasma membrane localization of Pit (Kawano *et al*., 2014a). The resting Pit is localized exclusively in the plasma membrane (Figure 7) (Kawano *et al*., 2014a), indicating that plasma membrane localization alone is not sufficient to trigger activation. The plasma membrane localization of Pit is ATP-binding activity-dependent because the P-loop mutant of Pit K203R is mislocalized (Kawano *et al*., 2010). This feature is similar to other CNLs, including RPM1, TM-22, and RPS5, whose auto active mutants are primarily localized to the plasma membrane (Qi *et al*., 2012; Chen *et al*., 2017; El Kasmi *et al*., 2017).

Recently, the structures of active and inactive forms of ZAR1 have been reported, revealing a more detailed structural observation of the NLR protein. Inactive ZAR1 forms a monomeric complex with resistance-related kinase (RKS1). *Xanthomonas campestris* pv. *campestris* AvrAC uridylates the PBS1-like protein 2 (PBL2) kinase to produce PBL2UMP, which triggers the pentameric ZAR1–RKS1–PBL2UMP resistosome *in vitro* and *in vivo* (Wang *et al*., 2019a; Wang *et al*., 2019b; Hu *et al*., 2020). Resistosome formation is required for AvrAC-triggered cell death and disease resistance. The *Pseudomonas syringae* effector HopZ1a also induces the oligomerization of ZAR1 *in vivo* (Hu *et al*., 2020). During the transition from the inactive to the active states of ZAR1, positional translation through unfolding and refolding in the a4 helix allows the α1 helix to be released from the four-helix bundle (Wang *et al*., 2019a). This conformational change of the α1 helix leads to the pentameric funnel-shaped structure of the CC domain of ZAR1. The funnel-shaped structure of active ZAR1 is similar to previously characterized pore-forming proteins, such as mixed lineage kinase-like (MLKL) and hemolytic actinoporin fragaceatoxin C (FraC) (Tanaka *et al*., 2015). Notably, FraC showed a similar conformational change during pore formation to that upon the activation of ZAR1. The N-terminal helix of FraC is released from the monomer and is capable of forming a funnel-shaped octamer, leading to its insertion into the cell membrane. The structure of the ZAR1 oligomer implies that the funnel structure of the CC domain of the ZAR1 oligomer also inserts into the cell membrane and induces cell death (Wang *et al*., 2019a). It appears that a funnel-shaped structure participates in membrane localization (Adachi *et al*., 2019). Recently, Adachi et al. found the consensus sequence called MADA motif in the N-termini of various CNLs which matches the N-terminal *α*1 helix of ZAR1. They predicted three residues mapped to the outer surface of the funnel-shaped structure of NRC4 based on the ZAR1 resistosome structure and substituted these three hydrophobic residues for negatively charged Glu residues. Those mutants failed to trigger cell death in *N. benthamiana* and one of the mutants decreased its plasma membrane localization, showing the general importance of insertion of *α*1 helix of CNLs into plasma membrane on their immunity. Since the membrane localization of Pit is also important for its function (Kawano *et al*., 2014a), it is possible that the CC domain of Pit has a funnel-shaped structure similar to that of active ZAR1 and plays an important role in its membrane localization and cell death. Our experiments showed that Pit I34E, L37E, and L41E mutants perturbed membrane localization and were localized in the cytoplasm (Figure 7A, B). In the active Pit model structure based on the active ZAR1 (PDB code 6J5T), the three hydrophobic residues (I34, L37, and L41) are located in the α2 helix but not in the α1 helix of the funnel-shaped structure, suggesting that the three hydrophobic residues are not directly involved in membrane insertion. The three hydrophobic residues, I34, L37, and L41, in the α2 helix of the Pit CC domain interact hydrophobically with V75, I78, and V79 in the α3 helix. The α3 helix is hydrophobically associated with I500 in the WHD domain and L510 in the LRR domain (Figure 6C). In addition, D77 in the α3 helix also forms a hydrogen bond with K532 in the LRR domain (Figure 6C). D77 is located at the EDVID motif in Pit (DDIVD in Pit) which is a highly conseved motif in CNLs (Bai *et al*., 2002). The EDVID motif directly contacts with LRR domain in the inactive ZAR1 structure (Burdett *et al*., 2019; Wang *et al*., 2019a). In the full-length MLA10 protein, the mutations of the EDVID motif in MLA10 weaken immune response but the same mutations in the CC domain fragment do not affect its autoactivity (Bai et al., 2012), suggesting that the EDVID motif is necessary for both autoinhibition and activation of MLA10. Like the ZAR1 case, it is possible that the EDVID motif serves as a signal relay from the LRR domain to the CC domain to indue the large conformational changes in the NB-LRR region. The three hydrophobic residues (I34, L37, and L41) in the α2 helix may support the funnel-shaped structure through interaction with the α3 helix, which is associated with the WHD and LRR domains. However, the substitutions of I34, L37, and L41 with Glu may destabilize this foundation for the funnel-shaped structure and consequently affect the insertion of the funnel-shaped structure formed by the N-terminal α1 helix into the membrane. Alternatively, we previously found that palmitoylation is required for plasma membrane localization of Pit (Kawano *et al*., 2014a) and these mutations in the CC domain of Pit may affect appropriate palmitoylation. But these speculations need to be tested in the future. The mislocalization of Pit by the mutations into the conserved hydrophobic residues disrupted the appropriate localization of the Pit-OsSPK1 complex (Figure 7C). This may lead to the attenuation of Pit-mediated immune responses.

## Author Contributions

Q. W. and Y. K. designed the study; Q. W., Y. L., K. K., J. L., D. Z., and Y. K. performed experiments and analyzed data; Q. W., K. K., and Y. K. wrote the manuscript; C. L. and D. M. gave technical support; Y. K. provided conceptual advice.

## EXPERIMENTAL PROCEDURES

### Plasmid Construction

For Gateway system-constructed plasmids, the target genes and fragments were first cloned into the pENTR/D-TOPO vector (Invitrogen), and then transferred by LR reaction into multiple destination vectors depending on the experimental requirements. For site-directed mutagenesis, overlapping PCR amplification using site-specific and mutagenic primers and pENTR templates was employed to generate *Pit* mutants. For pull-down and yeast two-hybrid assays, *Pit* mutants were directly cloned into the 6×His-SUMO, pGADT7, and pGBKT7 vectors, using restriction enzymes and T4 ligase (New England Biolabs).

### Yeast Two-Hybrid Assay

The Y2HGold-GAL4 system was used to test interactions between target proteins by transforming GAL4-AD/BD fused *Pit* plasmids into Y2HGold chemically competent cells (Weidibio: YC1002).

### Transient Expression and HR Assays

Agroinfiltration of *N. benthamiana* was conducted as described previously (Kawano *et al*., 2010). *Agrobacterium tumefaciens* strain GV3101 carrying the helper plasmid pSoup and binary plasmids was grown overnight at 28°C to an optical density at 600 nm (OD_600_) of around 0.8. Agrobacterial cells were harvested, resuspended in 10 mM MgCl_2_, 10 mM MES-NaOH (pH 5.6), and 150 µM acetosyringone, adjusted to OD_600_ = 0.4, and incubated at 23°C for 2–3 h before infiltration. We also used the p19 silencing suppressor to enhance gene expression. For coexpression of two proteins, *Agrobacterium* carrying the appropriate two constructs and p19 helper plasmid-containing bacteria were mixed at 1:1:1 volume ratio. The uppermost 3 or 4 leaves of 4-week-old *N. benthamiana* plants were selected for injection, and inoculated plants were kept in a growth room at 25°C for 2 days.

Transient expression of Pit mutants in tobacco leaves was performed according to the method described above. Each bacterial inoculum was infiltrated in a circle with a diameter of 1 cm on each of 15 leaves for three independent experiments. After 2–3 days, cell death symptoms became visible and were photographed.

For ROS detection and quantification assay, the infiltrated leaves expressing Pit mutants or negative control GFP were collected and floated in 1 mg/ml DAB solution for 5 h at room temperature. To visualize ROS *in situ*, the leaves were then decolorized with ethanol by boiling several times in a microwave oven until the chlorophyll was removed completely. ROS production of each sample was quantified by measuring the pixel intensities of the infected regions using ImageJ software (National Institutes of Health). The mean pixel intensity from three spots outside the infiltrated regions on each leaf was used to subtract background. Relative DAB staining intensity was calculated based on the mean pixel intensity of the GFP-infected region on each leaf to compare between different leaves.

### Plant Growth and Infection

All of transgenic rice plants used in this study were produced by the core facility of Shanghai Center for Plant Stress Biology. T0 generation of transgenic plants were used for infection analysis because introduced *Pit* WT gene did not sucessfully transmit T1 generation due to unknown reasons. Nipponbare plants were grown at 30°C for 5–6 weeks before being infected with the *M. oryzae* strain Ina86-137 (Race 007.0) (Hayashi & Yoshida, 2009). Infection of leaf blades by the punch method was performed as reported previously (Ono *et al*., 2001; Kawano *et al*., 2010). Lesion length and fungus growth were measured at 7 dpi. Photographs of disease lesions were taken at 6 dpi.

### Expression Analysis

Total RNA from rice was extracted using TRIzol reagent (Invitrogen). Total RNA (500 ng) was used for cDNA synthesis with a commercial kit (Vazyme) according to the manufacturer’s protocol. The cDNA was analyzed semi-quantitatively using normal polymerase mix. Total genomic DNA was extracted by the CTAB method and then subjected to quantitative analysis using SYBR Green Supermix (Bio-Rad) on a CFX96 Touch Real-Time PCR Detection System (Bio-Rad). *OsUbiquitin* was used as an internal control for normalization. Sequences of RT-PCR and RT-qPCR primers are listed in supplemental Table 1.

### Raichu-OsRac1 FRET Analysis

The Raichu intramolecular FRET system was applied as described previously (Kawano *et al*., 2010; Wong *et al*., 2018). Rice protoplasts were transformed with *Raichu-OsRac1* and *Pit* mutants or *GUS* vectors by the PEG method. Images of transformed cells were captured using a LEICA SMD FLCS microscope. Raichu-OsRac1 was excited using a 440 nm solid-state laser. The Venus and CFP filters were 550 ± 25 nm and 470 ± 20 nm, respectively.

### Protein Expression and Purification

His-SUMO tag- and glutathione-S-transferase (GST) tag-fused Pit CC (amino acids 1-140) were expressed in *Escherichia coli* strain BL21(DE3) Codon Plus. The bacteria were cultured at 37°C until the OD_600_ of the suspension of the medium was around 0.8. The recombinant proteins were induced with 0.3 mM IPTG for 12 h at 18°C. For protein purification, the bacterial cells were collected, resuspended, and sonicated in a lysis buffer (20 mM Tris-HCl [pH 8.0], 150 mM NaCl, 1 mM DTT). The proteins were then purified by affinity chromatography using Ni-NTA agarose resin and Glutathione Sepharose 4B resin (GE Healthcare), respectively.

### *In Vitro* Pull-down Assay

Equal amounts of His-SUMO-Pit CC and GST-Pit CC WT or mutated proteins in binding buffer (50 mM Tris-HCl [pH 7.5], 150 mM NaCl, 0.1% Triton X-100, 1 mM EDTA, and 1 mM DTT) were mixed to 200 μl for each reaction and incubated at 4°C for 5 min with gentle rotation. Glutathione Sepharose 4B resin was added to the solution for precipitation. The beads were washed five times with binding buffer and separated from the solution by centrifugation at 2500 × *g* for 2 min. The proteins were then eluted with 100 µl 2×SDS loading buffer for immunoblotting. Anti-SUMO (GenScript: A01693) and anti-GST (Abmart: M20007) antibodies were used.

### Subcellular Localization

Confocal fluorescence pictures were recorded under a Leica TCS-SP8 microscope, using 60×water-immersion objectives. A 488-nm laser was used to image GFP; a 514- nm laser was used to image Venus; and a 598-nm laser was used to image mCherry. For samples stained with a plasma membrane marker, FM4-64 (Invitrogen: F34653) solution was injected to the infiltrated leaves before harvesting and observation. The signals of FM4-64 were excited with a 566-nm laser.

### Luciferase Activity Assay in Rice Protoplasts

Isolation of rice protoplasts and PEG transformation were performed as described previously (Kawano *et al*., 2010; Wang *et al*., 2018). Protein preparation and luciferase– substrate interaction were conducted with a Luciferase Assay Report Kit (Promega). Luciferase activity was measured by a microplate reader (Thermo Scientific Varioskan Flash). We used mean values of three independent replications.

### Co-immunoprecipitation Assays

An *in vivo* co-immunoprecipitation (co-IP) assay was performed as previously reported (Wang *et al*., 2018). The infiltrated tobacco leaves were ground to powder in liquid nitrogen and homogenized in IP buffer (50 mM Tris-HCl [pH 7.5], 150 mM NaCl, 10% glycerol, 0.2% NP-40, 1 mM EDTA, 5 mM DTT, and EDTA-free protease inhibitor [Roche]). Anti-GFP agarose resin (Chromotek, GFP-Trap A, gta-20) was added to the extracted protein solution for precipitation. The GFP beads were washed five times and then subjected to immunoblot analysis together with input samples. Anti-Myc (Cell Signaling: #2276S), anti-HA (Roche: 11867423001), and anti-GFP (Abcam: ab6556) antibodies were used for western blot.

### Bimolecular Fluorescence Complementation (BiFC)

OsSPK1-cYFP, cYFP-OsRac1, and Pit-nYFP mutants were transiently expressed in *N. benthamiana* leaves according to the method described above. The signals of YFP were observed under a Leica TCS-SP8 microscope. A 514-nm laser was used to excite the YFP fluorescence and signals between 525 nm and 575 nm were recorded.

### Statistical Analysis and Biological Repetitions

Means were compared by using a *t* test (two-tailed; type 2). SEs were calculated in Microsoft Excel software. All of assays (including Co-IP, Y2H, Pull down, etc.) in this study were independently repeated at least three times.

## Supplemental information includes the following items

1. **Two supplemental figures and legends**. Figure S1 is related to Figure 3. Figure S2 is related to Figure 4.
2. **One supplemental table** Table S1. List of primers for experimental procedures

**Figure S1.**
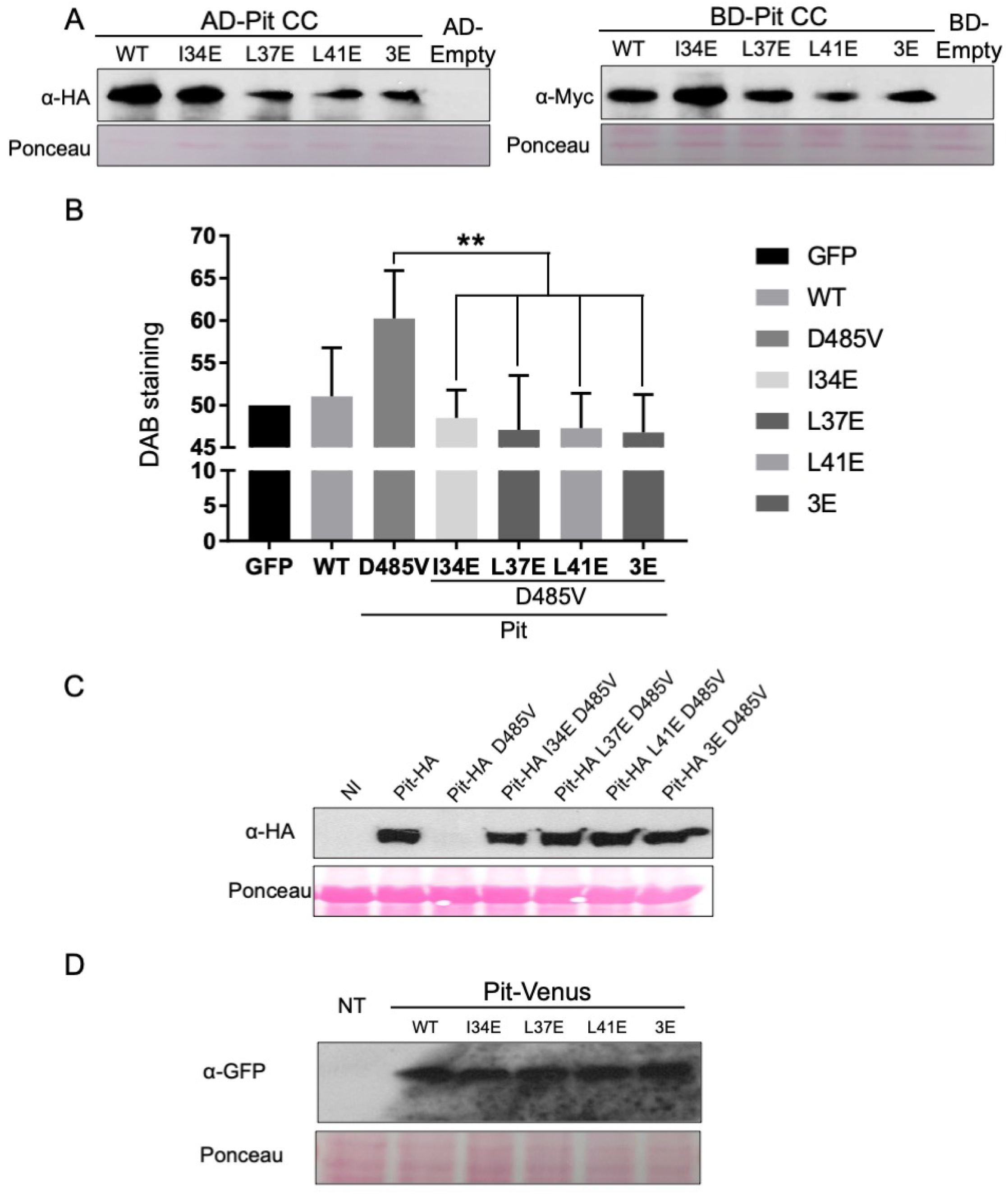
Effect of conserved hydrophobic residue mutations on Pit signaling. **A**, Protein expression of Pit CC WT and mutants in Y2HGold yeast cells. Anti-HA and anti-Myc antibodies were used for western blot to detect baits and preys, respectively. The post-transfer membrane was stained with Ponceau S. **B**, Quantitative analysis of the effect of I34, L37, and L41 mutation on Pit D485V-induced ROS production in *N. benthamiana*. Bars indicate DAB staining intensity relative to that observed after infiltration with negative control GFP. Data are expressed as mean ± standard error (SE) (**: *P* < 0.01; *n* = 10). Relative intensity of DAB staining (GUS=50) is shown. **C**, HA-tagged Pit mutants were transiently expressed in *N. benthamiana* leaves. After 2 days, the total proteins of infiltrated leaves were extracted for immunoblotting with anti-HA antibody. NI indicates non-infiltrated leaves. The post-transfer membrane was stained with Ponceau S and used as an internal control. **D**, Venus-tagged Pit mutants were transiently expressed in rice protoplasts. After 14 h, total protein was extracted with SDS loading buffer, and western blotting was then carried out with anti-GFP antibody. NT indicates a non-transformed sample. The post-transfer membrane was stained with Ponceau S.

**Figure S2.**
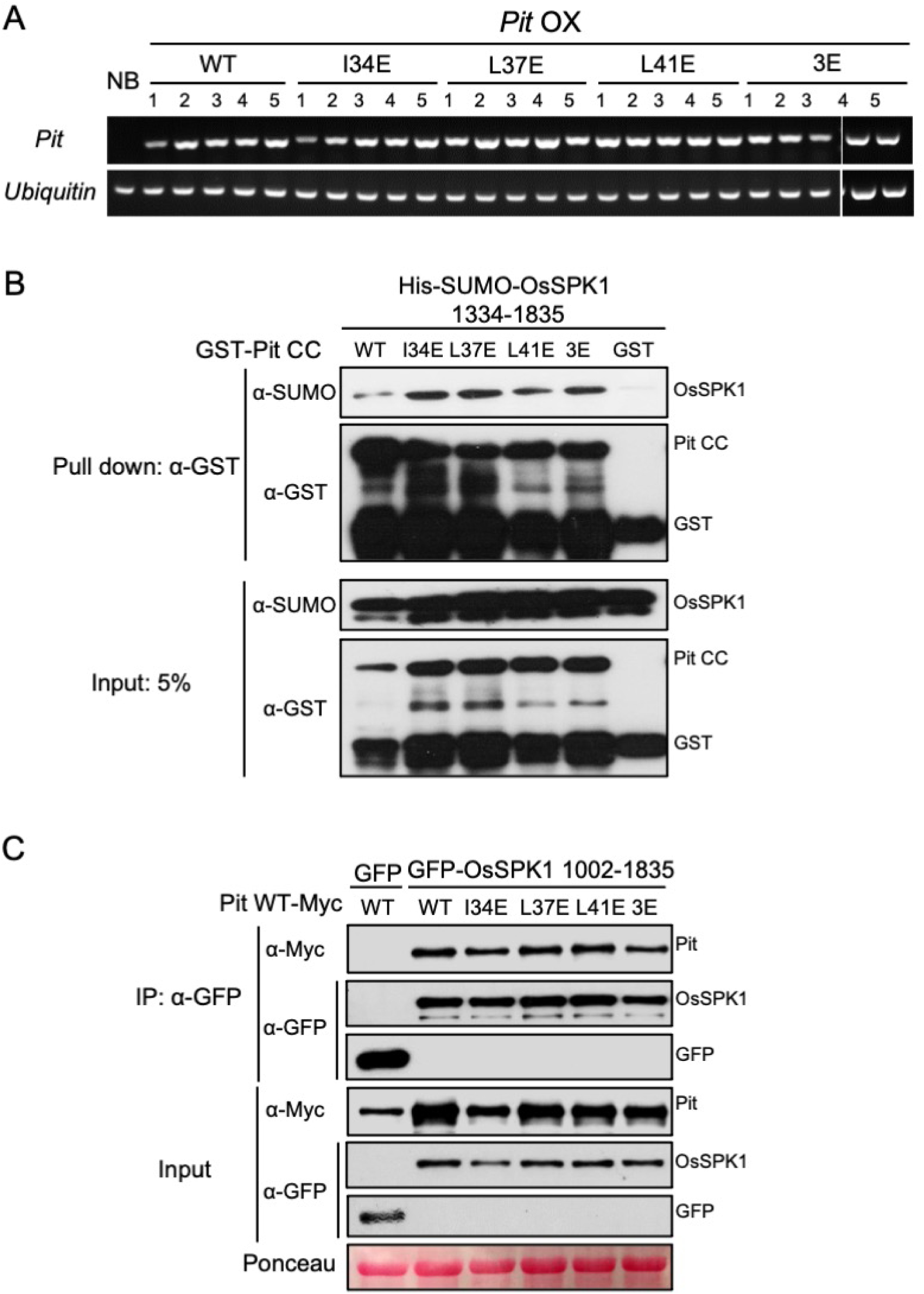
Three hydrophobic residues in the Pit CC domain are not necessary for binding to OsSPK1. **A**, Transcript levels of exogenous *Pit* WT and *Pit* mutants were measured by RT-PCR. Numbers indicate independent transgenic lines. *Ubiquitin* was used as an internal control. **B**, *In vitro* binding assay between Pit CC mutants and OsSPK1. Purified GST or GST-tagged Pit CC mutants immobilized on Sepharose were incubated with His-SUMO-tagged OsSPK1 (amino acids 1334–1835). After washing, the bound proteins were eluted by addition of SDS loading buffer. Anti-GST and anti-SUMO antibodies were used for subsequent western blotting analysis. **C**, Co-IP to analyze the interaction in *N. benthamiana* between OsSPK1 (amino acids 1002–1835) and full-length Pit with mutations in three conserved hydrophobic residues. Total protein extract was immunoprecipitated with anti-GFP antibody, and western blotting was then carried out with anti-GFP and anti-Myc antibodies. The post-transfer membrane was stained with Ponceau S.

**Table S1.**
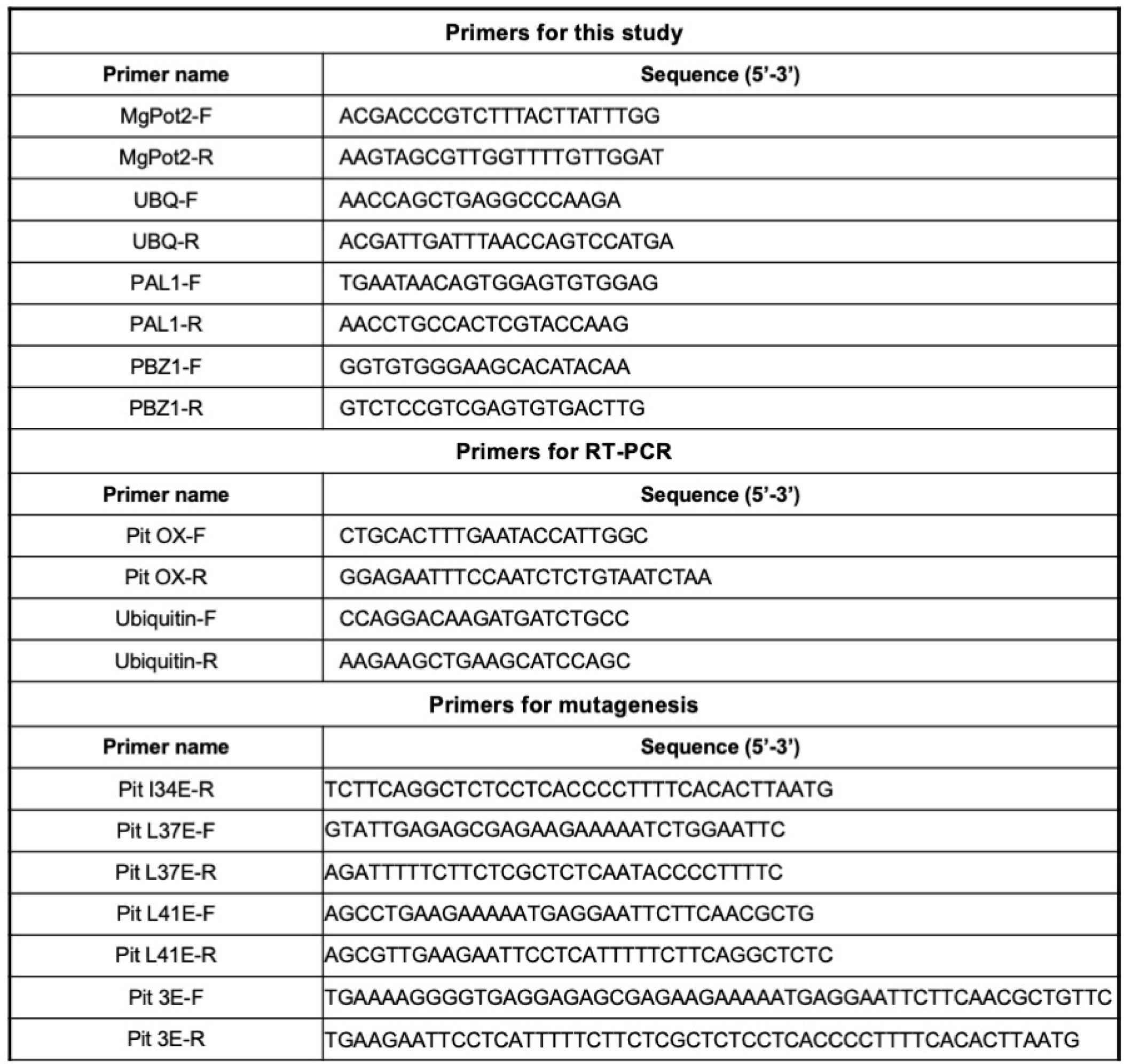

